# Short-Duration RAGE Antagonism Transiently Disrupts Tendon Homeostasis and does not Alter Diabetic Tendon Healing

**DOI:** 10.1101/2021.05.11.443619

**Authors:** Anne E. C. Nichols, Samantha N. Muscat, Alayna E. Loiselle

## Abstract

Obesity and type II Diabetes Mellitus (T2DM) have substantial pathological effects on tendon homeostasis, including loss of collagen organization and increased risk of tendon rupture. Moreover, following rupture or acute injury, the healing process is impaired by T2DM. We have previously demonstrating that restoring normal metabolic function in a murine model of obesity/ T2DM is insufficient to blunt or reverse the progression of diabetic tendinopathy, indicating the need for identification of novel therapeutic approaches to both maintain tendon homeostasis, and to improve the healing process. RAGE, the Receptor for Advanced Glycation Endproducts has been implicated as a key driver of several diabetic pathologies. We have demonstrated that pharmacological antagonism of RAGE is sufficient to partially improve tendon healing in non-diabetic animals. Therefore, in the current study we tested the efficacy of blunted RAGE signaling, via treatment with a RAGE Antagonist Peptide (RAP), to improve tendon healing in the context of T2DM. While our study did not find a beneficial effect of short-term RAP treatment on the healing process of T2DM mice, we did identify several important challenges brought about by this model of diet-induced obesity and T2DM. Both high fat (HFD) and low fat diet (LFD) feeding shifted the temporal molecular profile of healing compared to standard laboratory chow fed mice. Moreover, RAP treatment resulted in a transient disruption in homeostasis in the contralateral control tendons of both HFD and LFD mice, and this was due to a potential interaction with the systemic response to tendon injury as this response was not observed in HFD and LFD fed mice that did not undergo tendon repair surgery. Collectively, these data highlight the complications associated with models of diet induced obesity, and the lean control diets that should be considered in future studies.

## Introduction

Tendons are dense collagenous tissues that connect bone and muscle and facilitate many aspects of skeletal locomotion. Tendons are frequently injured due to either traumatic injuries, progressive degeneration or supraphysiological loading that results in tendon rupture. Following injury, which often requires surgical repair, tendons are prone to a scar-mediated healing response which bridges the tendon injury site with disorganized scar tissue that is mechanically inferior to native tendon. This lack of regenerative healing leaves the tendon prone to re-injury and with functional deficits. Despite the frequency of these injuries and the associated complications, the molecular processes that regulate scar-mediated healing are not well-defined. We have recently identified the small calcium binding protein S100a4 as a central regulator of fibrotic tendon healing ^1^, with the Receptor for Advanced Glycation Endproducts (RAGE) being the predominant receptor for S100a4 signaling. Indeed, inhibition of RAGE signaling using a RAGE antagonist peptide improved functional healing outcomes and did not impair reacquisition of mechanical properties ^1^. Importantly, scar-mediated tendon healing is further exacerbated by co-morbidities such as Type II Diabetes Mellitus (T2D). Type II Diabetes disrupts tendon homeostasis ^2–4^, substantially increases the risk of tendon rupture ^5, 6^ and pushes the healing tendon toward a more pronounced scar-mediated/fibrotic response ^7–10^. RAGE levels, which are largely dictated by ligand levels ^11^, are dramatically up-regulated in T2DM ^12^, and RAGE inhibition can reverse some pathological effects of T2DM ^13, 14^, suggesting the therapeutic potential of RAGE inhibition. Interestingly, Advanced Glycation Endproducts (AGEs), a primary ligand for RAGE, are increased in T2D ^15^. However, while AGEs are important in modifying tendon cell function ^16^, they do not alter tissue-level tendon mechanical properties ^17, 18^, suggesting the need to focus on other RAGE ligands as mediators of pathologic response to tendon healing in the context of T2D, and motivates a focus on RAGE signaling more broadly. Indeed, S100a4 is increased in several diabetic pathologies including diabetic retinopathy ^19^, and more recently S100a4 has been identified as a possible early bio-marker for insulin resistance ^20^. Therefore, we hypothesized that S100a4-RAGE signaling was increased in T2D tendon repairs relative to lean controls, and that inhibition of RAGE signaling, via systemic treatment with a RAGE Antagonist Peptide (RAP), would restore functional and morphological outcome parameters of healing of T2D tendons to the level of non-diabetic controls.

## Results

### RAGE and S100a4 expression are increased during diabetic tendon healing

We first examined changes in *S100a4* and RAGE (*Ager*) expression over the course of healing in HFD and LFD tendon repairs. A significant increase in *S100a4* mRNA expression was observed in HFD repairs at D7 (LFD: 2.6 ± 0.74; HFD: 8.7 ± 0.12, p=0.013), D14 (LFD: 56.23 ± 11.40; HFD: 74.74 ± 5.68, p=0.044), and D28 (LFD: 5.22 ± 1.28; HFD: 11.33 ± 4.3, p=0.038), relative to LFD repairs at the same time-points. No differences in *S100a4* expression were observed between groups at D3 or D21 (Fig. 1A). No differences in *Ager* (RAGE) mRNA expression were observed between HFD and LFD repairs at any time-point (Fig. 1C). However, a substantial increase in extent of RAGE staining was observed at D21 post-surgery in HFD repairs relative to LFD (Fig. 1C), indicating up-regulation of S100a4-RAGE signaling in obese/T2D tendon healing and suggesting inhibition of S100a4-RAGE signaling as a possible therapeutic target to improve diabetic tendon healing.

**Figure 1.**
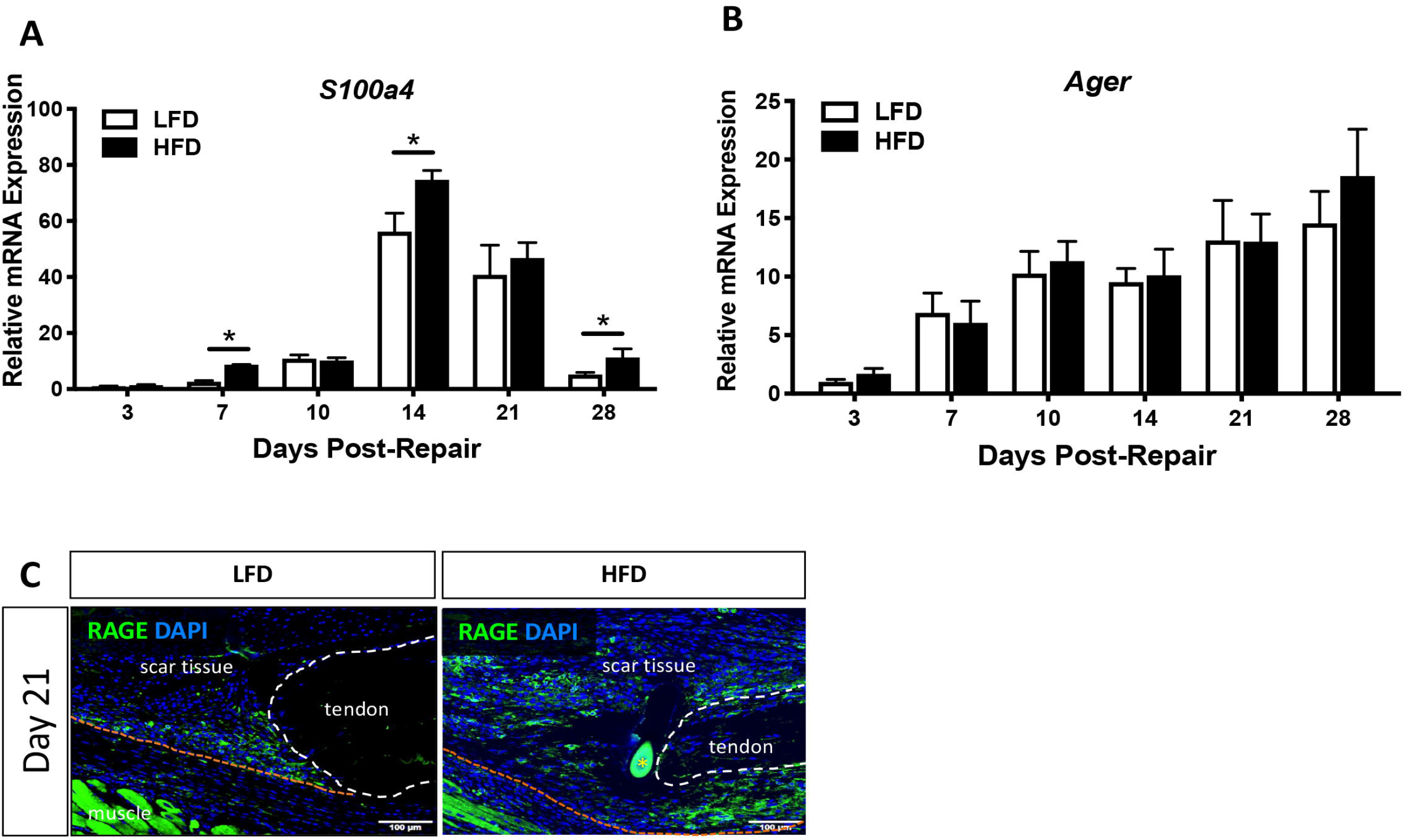
To examine changes in (A) *S100a4* and (B) *Ager* (RAGE) expression, mRNA was isolated from healing tendons of HFD and LFD-fed mice from 3-28 days post-surgery. (*) indicates p<0.05. (C) Immunofluorescence analysis of RAGE protein at D21 post-surgery demonstrates an expansion of RAGE expression in HFD-fed, relative to LFD-fed tendon repairs.

### RAGE antagonism transiently disrupts tendon homeostasis

To determine if inhibition of S100a4-RAGE signaling could improve obese/diabetic tendon healing, C57Bl/6J mice were placed on either a LFD or HFD beginning at 4 weeks of age, and were maintained on their respective diets for the duration of the experiment. Following tendon injury and repair surgery, mice were treated with RAGE Antagonist Peptide (RAP) or vehicle (0.5% bovine serum albumin [BSA]) from D5-10 post-surgery (Fig. 2A) based on the period of peak S100a4 expression^1^. Body weight of HFD-fed mice at harvest was significantly greater than that of LFD-fed mice at harvest at days 14, 21 and 28 post-surgery, regardless of drug treatment (Fig. 2B). There was no effect of RAP treatment on the body weight of mice on either diet at any timepoint (Fig. 2B).

**Figure 2.**
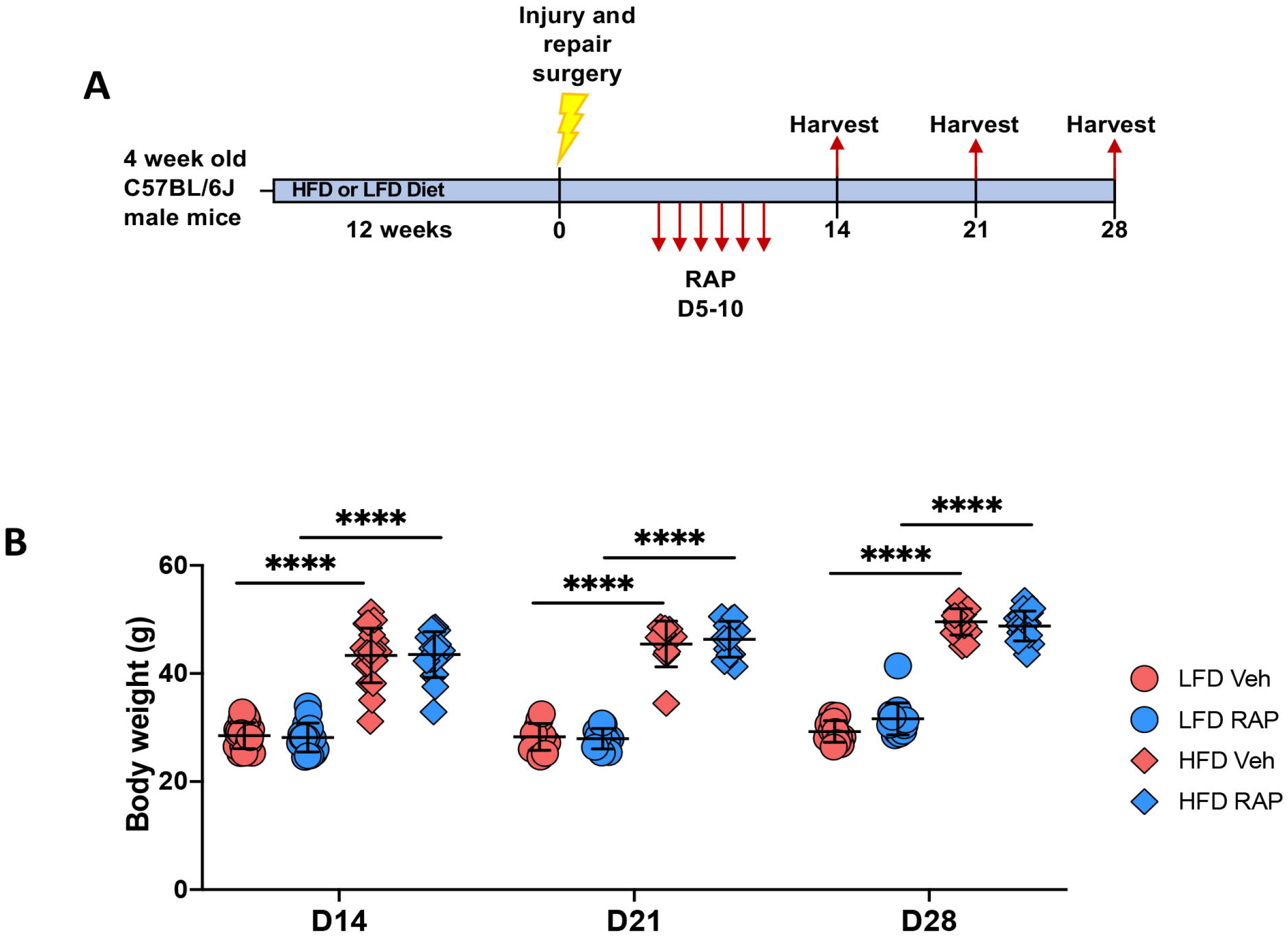
(A) Experimental design: 4-wk old male C57Bl/6J mice were initiated on either a High Fate Diet (HFD) or Low Fat Diet (LFD), and were maintained on their respective diets for the duration of the experiment. Mice were treated with RAGE Antagonist Peptide (RAP), or vehicle from D5-10 post-surgery. Tendons were harvested between 14-28 post-surgery. (B) Body weight at harvest from RAP and Vehicle treated HFD and LFD repairs. (****) indicates p<0.0001.

To determine if systemic RAP treatment has any effect on un-injured contralateral tendons, contralateral control hind paws were harvested at 14, 21 and 28 days post-surgery. Surprisingly, a significant decrease in max load at failure was observed in uninjured tendons from HFD and LFD RAP treated mice that were harvested at day 14 (LFD RAP: 10.03N ± 2.03 vs. LFD Veh: 12.66N ± 1.90, p=0.033; HFD RAP: 8.95N ± 2.2 vs. HFD Veh: 11.80N ± 2.20, p=0.026) relative to diet-matched vehicle treated mice (Fig. 3A). No differences in stiffness, MTP flexion angle or gliding resistance were observed as a function of treatment or diet (Fig. 3B-D).

**Figure 3.**
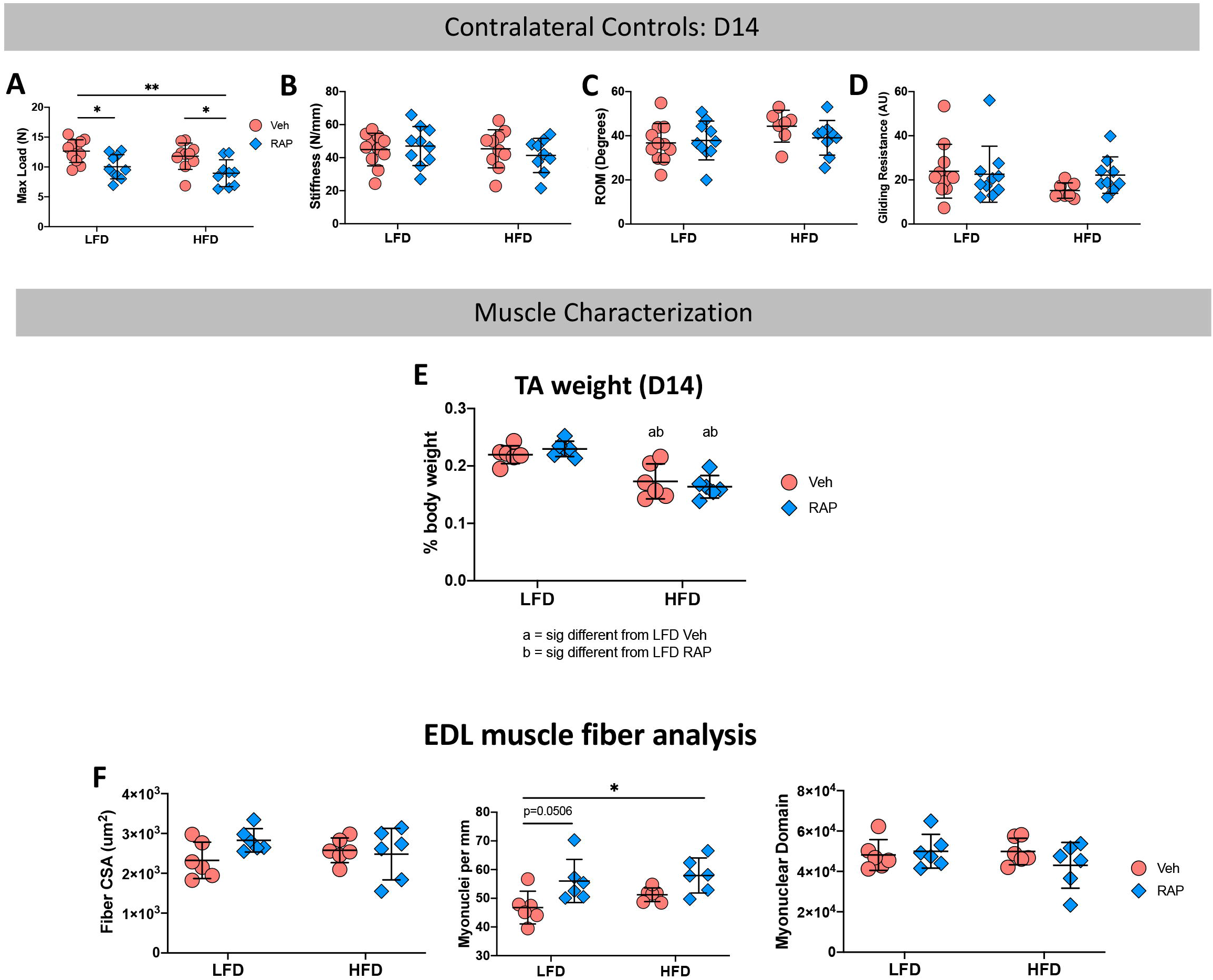
Contralateral (CL) control tendons harvested at day 14 post-surgery from RAP and Vehicle treated HFD and LFD mice underwent mechanical and functional testing to quantify (A) Max Load at Failure, (B) Stiffness, (C) Range of Motion (ROM), and (D) Gliding Resistance. To determine if alterations in CL tendon mechanical properties was due to an effect of RAP on the adjacent muscles, Tibialis Anterior (TA) and Extensor Digitorum Longus (EDL) muscles were harvested from day 14 CL limbs. (E) TA muscle weight, and (F) EDL Fiber CSA, myonuclear density, and myonuclear domain were quantified. (*) indicates p<0.05, (**) indicates p<0.01.

Based on this surprising result, we examined two potential reasons for this finding. First, to determine if systemic RAP treatment may alter muscle function which may impact tendon loading and therefore mechanical properties, we harvested tibialis anterior (TA) and extensor digitorum longus (EDL) muscles from the contralateral control limbs of mice on both diets 14 days post-repair. No significant differences in relative TA weight were observed between vehicle and RAP treated animals in either diet (LFD Veh: 0.22 ± 0.02% vs. LFD RAP: 0.23 ± 0.01%; p=0.833, HFD Veh: 0.17 ± 0.03% vs. HFD RAP: 0.16 ± 0.02%; p=0.864) though relative TA weight was significantly reduced in HFD mice compared to LFD mice regardless of RAP treatment (Fig. 3E). EDL muscles were used for muscle fiber characterization. No differences in fiber cross-sectional area were observed as a function of diet or treatment (Fig. 3F), however RAP treatment resulted in increased myonuclear density relative to vehicle in the LFD group (p=0.050). Myonuclear density was not different between RAP and vehicle-treated HFD muscles, though it was significantly increased in RAP treated HFD EDLs, relative to vehicle treated LFD EDLs (p=0.0149) (Fig. 3G). No differences in myonuclear domain were observed as a function of treatment or diet (Fig. 3H). Collectively these data demonstrate that RAP treatment may have modest systemic effects on some aspects of muscle function but is likely not the cause of decreased max at load at failure in RAP treated contralateral control tendons.

Second, to determine if the decrease in max load in D14 RAP treated contralateral control tendons was due to an interaction between RAP and tendon injury and a systemic response to tendon repair surgery, we also examined the effects of RAP treatment on HFD and LFD-fed naïve mice that did not undergo tendon repair surgery. No differences in max load at failure were observed between tendons from vehicle and RAP treated naïve mice in either the LFD or HFD group. However, max load at failure was significantly increased in tendons from RAP treated naïve HFD (p=0.305) and LFD mice (p=0.014), relative to RAP treated HFD contralateral tendons (Supplemental Table 1), suggesting a potential interaction between RAP treatment and tendon surgery.

At D21 post-surgery, no differences in max load at failure, stiffness, MTP Flexion angle, or Gliding resistance (Supplemental Table 2) were observed in contralateral control tendons as a function of either HFD/LFD or treatment. Consistent with this, no differences in max load at failure, stiffness, MTP Flexion Angle, or Gliding Resistance were observed between contralateral control tendons of HFD and LFD-fed mice treated with RAP or vehicle at D28 (Supplemental Table 2). Collectively, these data suggest that systemic RAP antagonism transiently disrupts tendon homeostasis soon after treatment, with these effects diminishing with time, and also resulting from an interaction between RAP and the systemic response to tendon injury.

We also sought to understand if the high-fat and low-fat diets altered the response to RAP, relative to standard lab chow, which has been used in our prior RAP study ^1^. At D21 post-surgery, HFD and LFD CL-controls were not significantly different than normal chow-fed CL-controls in terms of max load at failure, MTP flexion angle, or gliding resistance. However, significant differences in stiffness were observed between LFD Vehicle (p<0.01), HFD vehicle (p<0.01) and HFD RAP-treated tendons (p<0.01), compared to normal-chow-fed RAP treated tendons (Supplemental Table 3). At D28, no differences in max load or stiffness were observed in HFD and LFD CLs, relative to normal-chow-fed CLs. However, a significant increase in MTP Flexion Angle was observed in LFD RAP CLs compared to normal chow RAP-treated CLs (p=0.049), and an increase in MTP Flexion Angle was observed in HFD RAP CLs, relative to normal chow RAP CLs; however, this did not reach statistical significance (p=0.051). Consistent with this, gliding resistance was decreased (though not significantly) in LFD RAP CLs (p=0.0718) and HFD RAP CLs (p=0.0706), compared to normal chow RAP CLs (Supplemental Table 3). Taken together, these data highlight the important interaction between diet composition, systemic response to injury and efficacy of pharmacological antagonism of RAGE.

### RAGE Antagonism does not alter tendon healing in HFD- and LFD-fed mice

At D14 post-surgery, no differences in MTP ROM, gliding resistance, max load at failure, or stiffness (Fig. 4A), as a function of either diet of RAP treatment. Consistent with this, no changes in any functional parameters were observed due to either diet or RAP treatment at D21 (Fig. 4B), or D28 (Fig. 4C).

**Figure 4.**
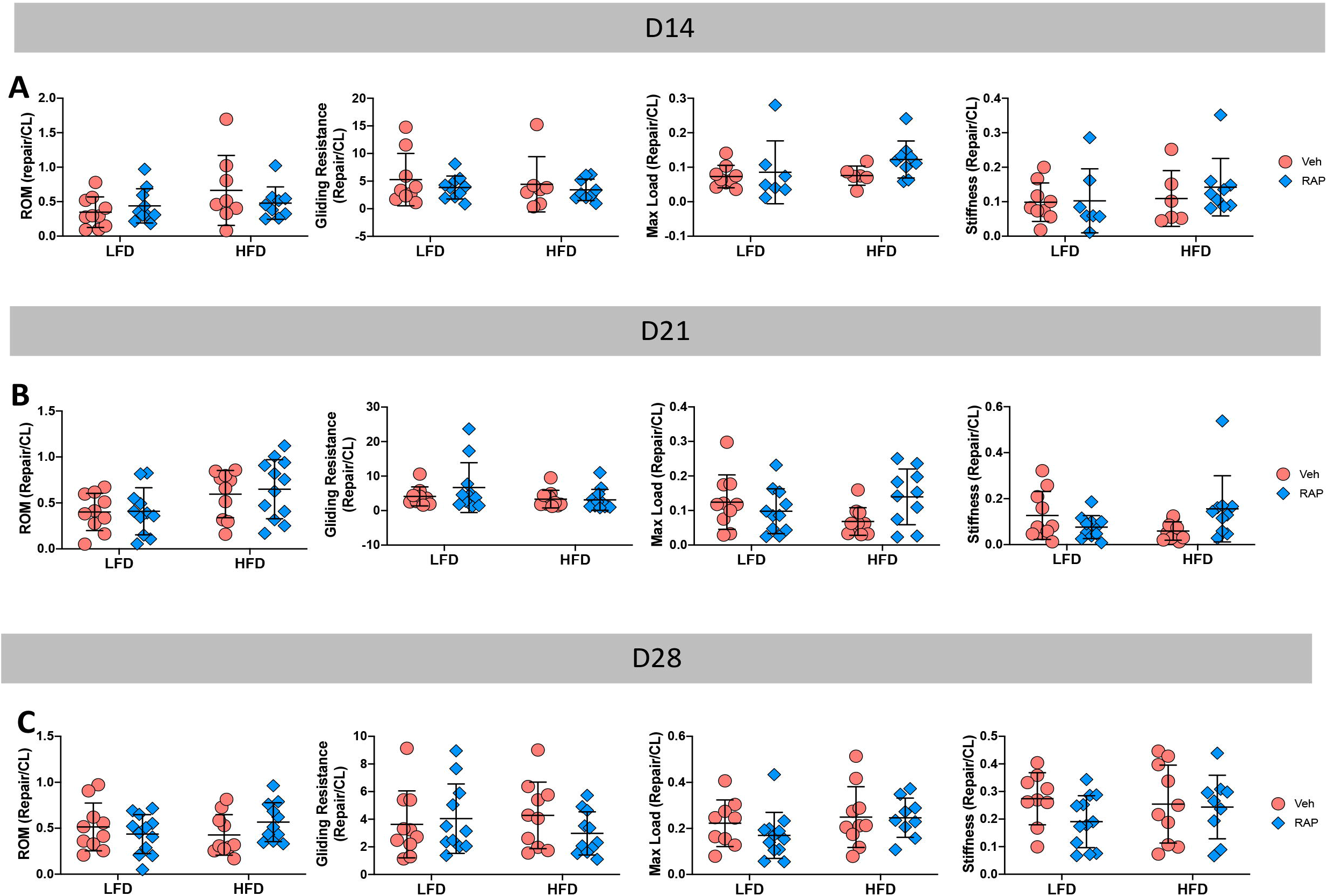
To examine the effect of RAP on the healing process, tendons were harvested at (A) D14, (B) D21, (C) D28 post-surgery, and Range of motion (ROM), Gliding Resistance, Max load at failure, and stiffness were quantified.

To better understand the surprising lack of effect of both HFD and RAP treatment on the healing process, we assessed both total RAGE expression and signaling pathways down stream of RAGE. At D14 post-surgery, no differences in RAGE expression were observed between groups (Fig. 5A). However, a substantial decrease in phospho-p65, an indicator of NFκB activation was observed in LFD-RAP, relative to LFD-Veh, suggesting successful RAGE antagonism (Fig. 5B). Surprisingly, no differences in p-p65 were observed between HFD-RAP and HFD-Veh (Fig. 5B). No changes in pERK1/2 were observed between groups (Fig. 5B).

**Figure 5.**
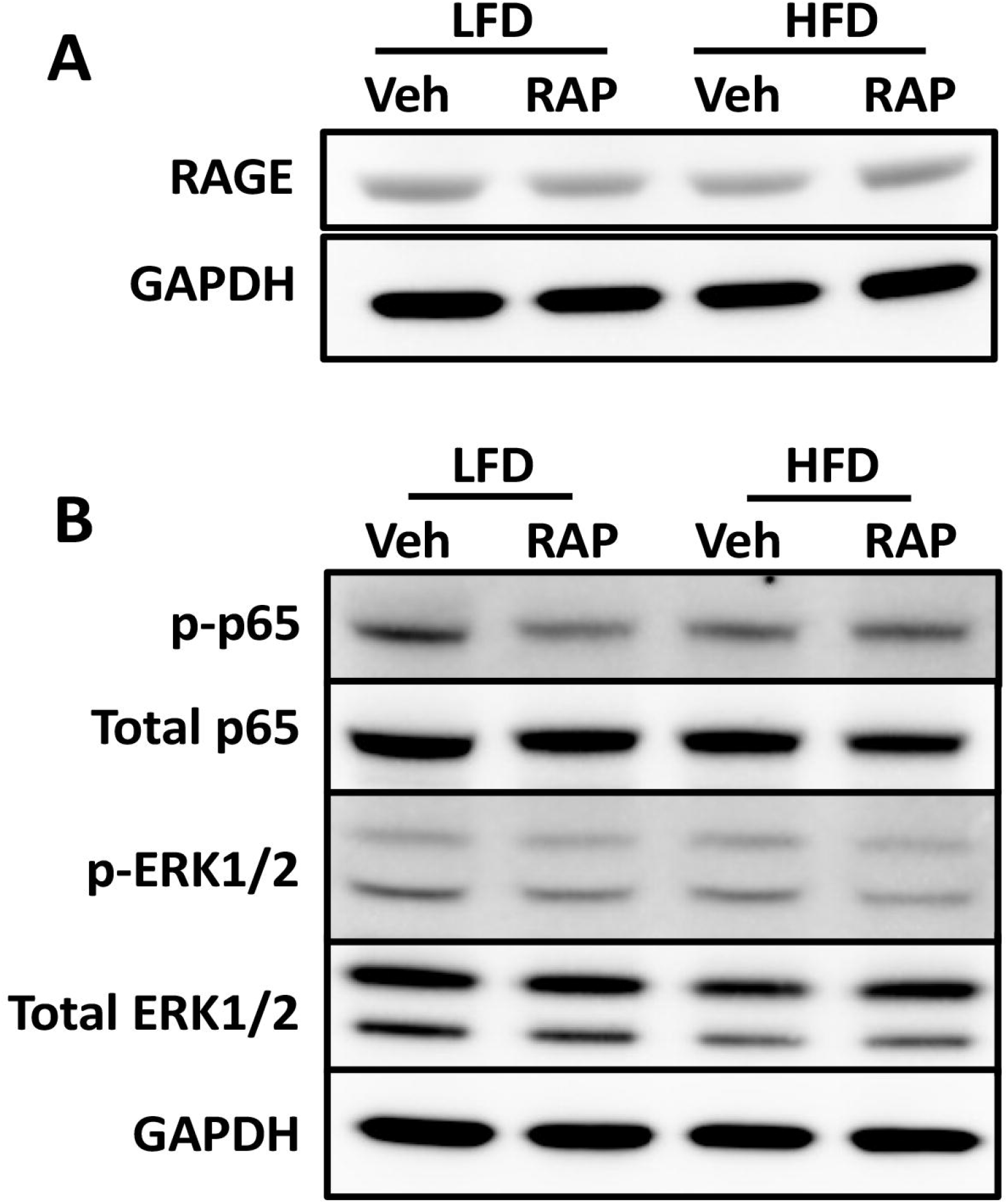
Western blot assessment of (A) total RAGE protein, normalized to GAPDH, and (B) activation of NF B signaling, as indicated by phospho-p65 levels, which are normalized to total p65 levels, and pERK1/2 activation, normalized to total ERK1/2 expression at D14 post-surgery in HFD and LFD-fed RAP and vehicle treated tendons.

## Methods

### Mice and model of T2DM/obesity

Male C57BL/6J mice (#664, Jackson Laboratories) were placed on a HFD (60% Kcal from fat; #D12492; Research Diets, Inc, New Brunswick, NJ) for 12 weeks beginning at 4 weeks of age, which results in obesity and elevated blood glucose ^7, 21^. Lean control mice were fed a sucrose-matched LFD (10% Kcal from fat; #D12450J; Research Diets, Inc). Mice were maintained on their respective diets for the duration of the experiment, and body weight was recorded at the time of harvest. Only male mice were used in this study, as female C57Bl/6J mice fed a HFD become obese but not diabetic ^22^.

Given that the LFD is different than the standard laboratory chow (Laboratory Autoclavable Rodent Diet 5010, LabDiet, St. Louis, MO) that we have used in prior studies related to RAP ^1^, an additional cohort of mice (“chow-fed”) were fed standard lab chow for the duration of the experiment to further define the effects of diet modification on the tendon healing process and response to RAP treatment.

### Tendon injury and repair surgery

At 16-weeks of age, mice underwent surgical injury and repair of the flexor digitorum longus (FDL) tendon as previously described ^23–25^. Briefly, mice received sustained-release buprenorphine (15 to 20 μg) prior to surgery. Mice were anesthetized with ketamine (100 mg/kg) and xylazine (10 mg/kg). The FDL tendon was first transected at the myotendinous junction to protect the repair site, and the skin was sutured closed with 5-0 suture. A small incision was then made through the skin on the plantar surface of the right hindpaw to allow for location of the FDL tendon, which was then completely transected and repaired using 8-0 non-resorbable nylon suture. The skin over the repair was closed using 5-0 suture. Mice were closely monitored following surgery to ensure they regained normal ambulation and food and water intake.

Un-injured contralateral tendons were harvested as a control (“contralateral”; “CL”), however, to assess the potential impact of any systemic response to the tendon repair surgery, and how this may alter the RAP-treatment response, an additional cohort of ‘naïve’ animals were used, in which the tendon repair surgery was not performed.

### RAP treatment

To inhibit RAGE signaling, mice were treated with RAGE Antagonist Peptide (RAP), as we have previously described ^26^. Briefly, mice were treated with 100μg RAP (#553031, MilliporeSigma), or vehicle (0.5% bovine serum albumin in saline) via i.p. injection on D5-10 post-surgery.

### Immunofluorescence

Following harvest, tendons were fixed (10% neutral buffered formalin) for 72hrs, decalcified (14% EDTA for 14 days), and processed for routine paraffin histology. Three-micron sections were cut through the sagittal plane of the hindpaw and mounted on Superfrost Plus slides (VWR, Radnor, PA). For immunofluorescence analysis, sections were dewaxed, rehydrated, blocked in 10% normal donkey serum, and incubated in primary antibody overnight at 4°C (RAGE: 1:100, # sc-365154, SantaCruz). Following overnight incubation, sections were then washed and incubated with fluorescently-conjugated secondary antibody for 1hr at room temperature (RAGE: Goat anti-mouse AlexaFluor488 secondary; 1:1000; Cat. #: A11029, ThermoFisher). Sections were then stained with the nuclear dye DAPI, cover slipped, and imaged on an Olympus VS120 Virtual Slide Microscope.

### RNA extraction and Qpcr

Tendons were harvested between 3-28 days post-surgery from HFD and LFD-fed mice, and total RNA was extracted as previously described ^23, 26^. Briefly, tendons were homogenized in TRIzol reagent, and RNA was extracted according to manufacturer protocol. 500ng of mRNA was reversed transcribed into cDNA using iScript cDNA synthesis kit (Bio-Rad, Hercules, CA). Quantitative PCR (qPCR) was run using gene specific primers for *S100a4, Ager* (Rage), and *Gapdh* were used. Expression levels were normalized to *Gapdh*, and to expression in D3 LFD repairs. RNA was extracted from 3 tendons per diet per time-point.

### Western blotting

Total protein was extracted from HFD and LFD FDL tendons at 14 days post-repair. Samples contained the scar tissue as well as 1 to 2 mm of native tendon on either side of the repair site. Only intact repairs were used for protein and samples were pooled for each group (LFD Veh n=3, LFD RAP n=4, HFD Veh n=2, HFD RAP n=4) to ensure adequate protein yield. Tendons were homogenized in radioimmunoprecipitation assay (RIPA) buffer with protease/phosphatase inhibitors (Cell Signaling Technology [CST] #58725) using 0.5-mm zirconium oxide beads and a Bullet Blender Gold Cell Disrupter (Next Advance Inc.). Samples (35 μg) were then separated using NuPAGE 4 to 12% Bis-Tris Gels (Invitrogen #NP0335) and transferred to nitrocellulose membranes (Invitrogen #IB23011). Membranes were probed overnight at 4°C with antibodies to RAGE (1:1000; Santa Cruz #sc-365154), phospho-p65 (1:1000; CST #3033), total p65 (1:1000; CST #8242), phospho-ERK1/2 (1:1000; CST #4377), total ERK1/2 (1:1000; CST #9102), and glyceraldehyde-3-phosphate dehydrogenase (GAPDH) (1:1000; CST #2118) in 5% bovine serum albumin (total protein antibodies) or 5% dry milk (phospho-specific antibodies) Tris buffered saline (Bio-Rad #1706435) with 0.1% Tween-20. Bands were visualized using SuperSignal West PLUS Chemiluminescent Substrate Pico (total proteins; Thermo Scientific #34580) or Femto (Thermo Scientific #34095) and imaged on a ChemiDoc™ Imaging System (Bio-Rad). Band intensity quantification was performed using ImageLab (Bio-Rad, Version 6.0.1, build 34).

### Functional and biomechanical outcome assessment

Tendon functional outcomes were assessed as previously described ^23, 25–28^. Briefly, hindlimbs were dissected at the knee joint and the proximal end of the FDL tendon was isolated at the myotendinous junction. The end of the FDL tendon was then sandwiched between two pieces of tape which was used to incrementally load the tendon with weights ranging from 0 to 19 g.

Images were taken after at each load and were subsequently used to calculate the flexion angle of the metatarsophalangeal (MTP) joint using ImageJ. Range of motion (ROM) was calculated as the maximum MTP flexion angle at 19g load relative to the unloaded angle. Gliding resistance was determined from changes in the MTP flexion angle over the range of applied loads. After functional testing, biomechanical testing was performed. The FDL tendon was released from the tarsal tunnel, and the proximal end of the tendon and the toes of the hindpaw were gripped into an Instron 8841 uniaxial testing system (Instron Corporation). Tendons then underwent load to failure testing at a rate of 30 mm/min.

### Skeletal muscle analysi

At 14 days post-repair, the contralateral limbs, along with the tibialis anterior (TA) muscle were harvested from mice in each experimental group. TA muscles were immediately weighed to determine any anabolic or catabolic effect of systemic RAP treatment on skeletal muscle. Hindlimbs from the knee down were fixed for 48 hours in 4% paraformaldehyde. After rinsing, extensor digitorum longus (EDL) muscles were carefully dissected and placed into 40% sodium hydroxide for two hours to facilitate myofiber dissociation. Individual myofibers were washed in PBS and stained with DAPI to visualize myonuclei. Whole myofibers were then imaged (Olympus VS120 Virtual Slide Microscope) used to determine myofiber cross-sectional area (CSA), number of myonuclei, and myonuclear domain as described in ^29^. Twenty individual myofibers were counted and averaged for each mouse (n=6 mice per group).

### Statistical analysis

All statistical analyses were performed using GraphPad Prism (version 8.3.1). All data are shown as the mean ± standard deviation. P-values of ≤0.05 were considered statistically significant, with significance differences displayed on figures as follows: *p≤0.05, **p≤0.01, ***p≤0.001, and ****p≤0.0001.

## Discussion

Given the naturally fibrotic response to acute tendon injury, and the further decrements in the healing process that occur due to Type II Diabetes, there is a clear need to define translational therapies to enhance the healing process. We have previously shown that inhibition of S100a4 signaling promotes more regenerative healing ^26^, with RAGE being a key receptor to mediate downstream signaling. Thus, identifying S100a4-RAGE signaling as a potential therapeutic target. However, RAGE can also bind other ligands, including Advanced Therefore, we tested the efficacy of small-molecule inhibition of RAGE, via treatment with RAGE Antagonist Peptide (RAP), using a dose and time-course we have demonstrated to be efficacious in enhancing tendon healing in non-diabetic mice. Surprisingly, RAP had no effect on the healing process, although a transient disruption in homeostasis was observed in uninjured contralateral control tendons of both HFD and LFD-fed mice shortly after RAP treatment. Moreover, we also demonstrate that these effects are likely due to an interaction between RAP treatment and a systemic response to tendon injury, as these effects were not observed in RAP treated mice that had never undergone tendon surgery. Finally, we show that while LFD is an appropriate diet-matched control for HFD, the tendon healing response, and RAP responsiveness in these mice differs from non-diabetic mice that are fed standard laboratory chow. While this study did not support the efficacy of this specific RAP treatment paradigm, these data shed important light on many key variables and interactions that can impact both the healing process (e.g. diet composition, systemic response to injury), as well as the application of pharmacological candidates to the paradigm of diabetic tendon healing (e.g. changes in the temporal-molecular profile, drug efficacy).

The RAGE receptor has been widely studied as a central regulator of several diabetes-related pathologies, including nephropathy ^30^, retinopathy ^31^, and diabetic atherosclerosis ^14^. Interestingly, systemic deletion of RAGE, using RAGE-/-animals, protects animals from the development of diet-induced-obesity ^32^, further supporting a requirement for RAGE in these pathologies. Moreover, pharmacological inhibition using RAP, or blockade of RAGE signaling blunted the development of diabetic pathologies including atherosclerosis ^13, 14^ These studies, in combination with our previous work demonstrating that RAP improves range of motion following tendon injury ^26^, suggest RAGE as a potential central regulator impaired diabetic tendon healing. However, the present study did not support the efficacy of short-duration RAGE antagonism to improve tendon healing, though this is likely related to the temporal shift in RAGE expression in both HFD and LFD-fed mice, relative to that observed in standard laboratory chow fed animals ^26^. Despite this, we also observed several interesting effects of RAP treatment. Perhaps most notably is the transient impact of RAP treatment on the mechanical properties of contralateral control tendons. The lack of effect of RAP treatment on mice that had never undergone tendon repair surgery suggest that RAP treatment may interact with a systemic response to tendon injury. This is particularly interesting as the response to tendon injury is thought to be quite localized. Indeed, in prior studies that examined the recruitment of bone marrow derived cells to the tendon injury site, a robust recruitment was observed, and a complete lack of recruitment was observed in the contralateral control ^33^. While we did not assess this, these data may suggest a systemic increase in RAGE in response to injury, resulting in this surprising interaction between RAP and contralateral control tendon response.

There are several limitations to this study. First, the dose and timing of RAP treatment that was used is based on efficacy in normal chow fed mice ^26^. However, our data clearly demonstrate that both HFD and LFD alters the healing course relative to normal chow, including delaying the peak expression of RAP until later in healing. As such, we see limited efficacy of RAP treatment, likely to due to the fact that we are not antagonizing RAGE signaling at the optimal time. In addition, HFD and LFD animals were given the same dose of RAP, despite substantial differences in body weight. Future studies that modulate both the RAP treatment timing, as well as the dose on a per weight basis will be needed to fully establish the potential of RAP to enhance tendon healing. Finally, there are limitations related to this model of diet induced obesity. While we and others ^7–9, 34, 35^ have previously demonstrated detrimental effects of obesity/diabetes on the tendon healing process, minimal effects were observed in this study. We hypothesize that this may be due to variability of the degree of obesity/diabetes using this model. For example, in co-housed male mice a clear hierarchy is often established ^36, 37^, which may lead to variation in diabetic phenotype and therefore healing capacity, such that differences between HFD and LFD are not detected. Future work using genetic models of obesity (e.g. Lep^ob/ob 38^, Lepr^db/db 39^) may be useful in providing a more uniform healing response.

Type II Diabetes is a major risk factor for diabetic tendinopathy development, leading to a greater risk of tendon rupture. Moreover, the healing process is substantially impaired in both diabetic humans and pre-clinical animal models. Importantly, following the development of diabetic tendinopathy, restoration of normal metabolic function is insufficient to blunt or reverse tendon pathology ^21^, suggesting that alternative treatment approaches are needed. Therefore, we leveraged our previous success using a RAGE Antagonist Peptide to enhance tendon healing to determine the potential of this approach to improve diabetic tendon healing. However, this study highlights several challenges related to the use of modified diets, and the consistency of the diet-induced obesity model. Addressing these model limitations will be an important consideration for future work, which will ultimately enable identification of therapeutic strategies to maintain tendon function and improve tendon healing in diabetic patients.

## Acknowledgements

We would like to thank the Histology, Biochemistry and Molecular Imaging (HBMI) and the Biomechanics, Biomaterials and Multimodal Tissue Imaging (BBMTI) at the University of Rochester Medical Center for technical assistance with the histology and biomechanical testing, respectively. This work was supported in part by NIH/NIAMS T32 AR076950 (to AECN), K01AR068386 and R01AR073169 (to AEL). The HBMI and BBMTI Cores were supported by NIH/NIAMS P30AR069655. The content is solely the responsibility of the authors and does not necessarily represent the official views of the National Institutes of Health.

## Supplemental Tables

**Supplemental Table 1**. To determine if there was a systemic response to injury that interacted with the response to RAP treatment, an additional cohort of HFD and LFD mice were used in which mice were treated with RAP, or vehicle, but did not undergo tendon repair surgery. Range of motion, Gliding Resistance, Max load at failure and stiffness were quantified in naïve HFD and LFD controls four days after the final RAP/Vehicle treatment to match the D14 time-point. Statistical comparisons are shown between naïve groups, and compared to HFD and LFD, RAP and Vehicle treated CL control tendons harvested at day 14 post-surgery.

**Supplemental Table 2**. Summary of functional and mechanical data from day 21 and day 28 contralateral control tendons from HFD and LFD-fed mice treated with RAP or vehicle, and statistical comparisons to mice fed standard laboratory chow (“chow”) mice that were treated with RAP or vehicle and harvested at the matched timepoint. (a) indicates p<0.05 vs. Chow-fed vehicle treated contralateral control tendons (b) indicates p<0.05 vs. chow-fed RAP treated contralateral control tendons, (c) indicates p<0.05 vs. LFD-fed vehicle treated contralateral tendons.

**Supplemental Table 3**. Summary of functional and mechanical data from day 21 and day 28 post-surgery tendon repairs from mice fed a standard laboratory chow diet (“chow”), and treated with RAP or vehicle, and statistical comparisons to HFD and LFD-fed mice that were treated with RAP or vehicle and harvested at the matched timepoint. (a) indicates p<0.05 vs. HFD-fed vehicle treated tendon repairs, (b) indicates p<0.05 vs. LFD-fed vehicle treated tendon repairs, (c) indicates p<0.05 vs. HFD-fed RAP treated tendon repairs, (d) indicates p<0.05 vs. LFD-fed RAP treated tendon repairs.

