## Supplementary figures and images for "Short-Duration RAGE Antagonism Transiently Disrupts Tendon Homeostasis and does not Alter Diabetic Tendon Healing"

### Supplemental Table 1

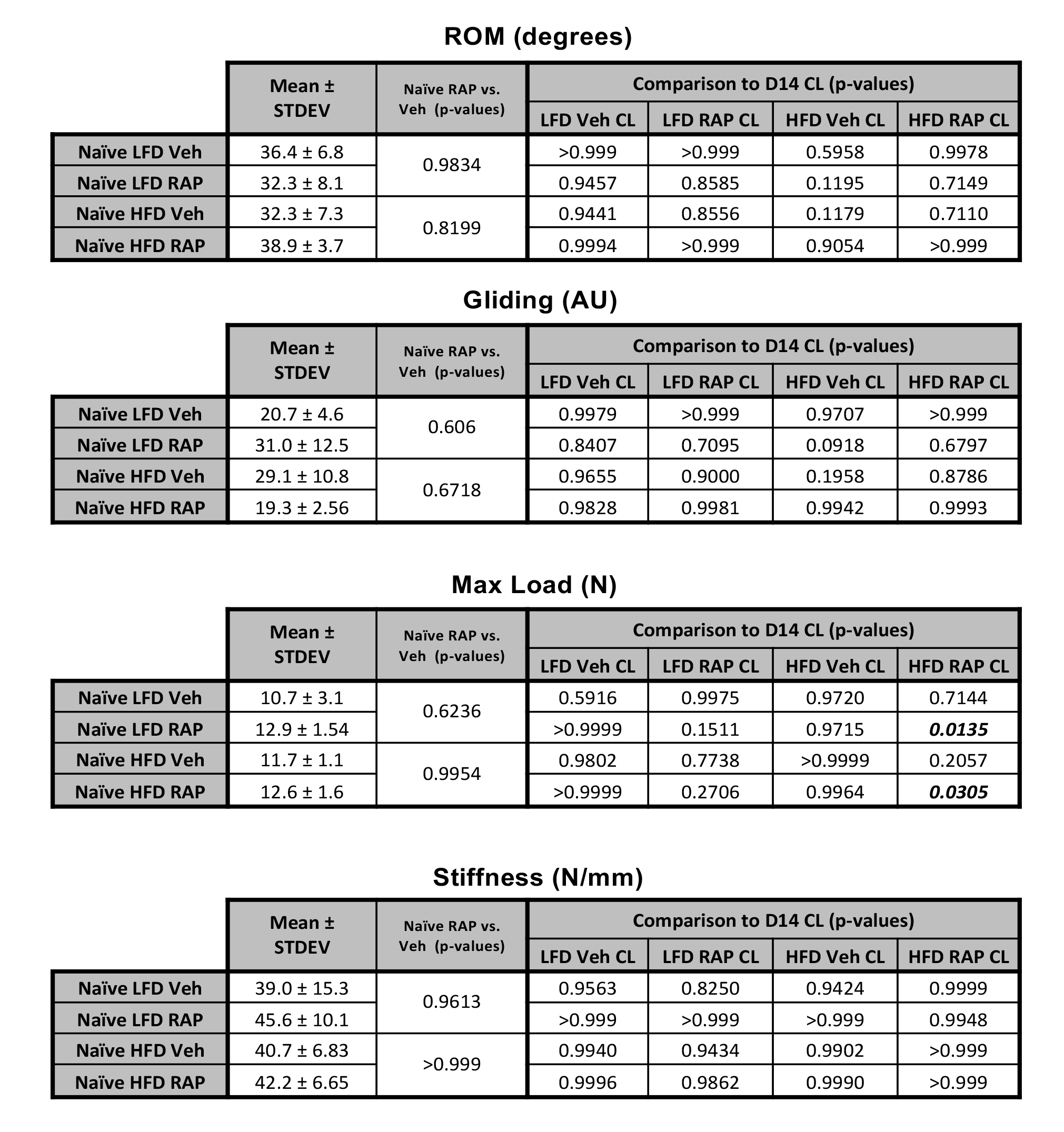

### Supplemental Table 2

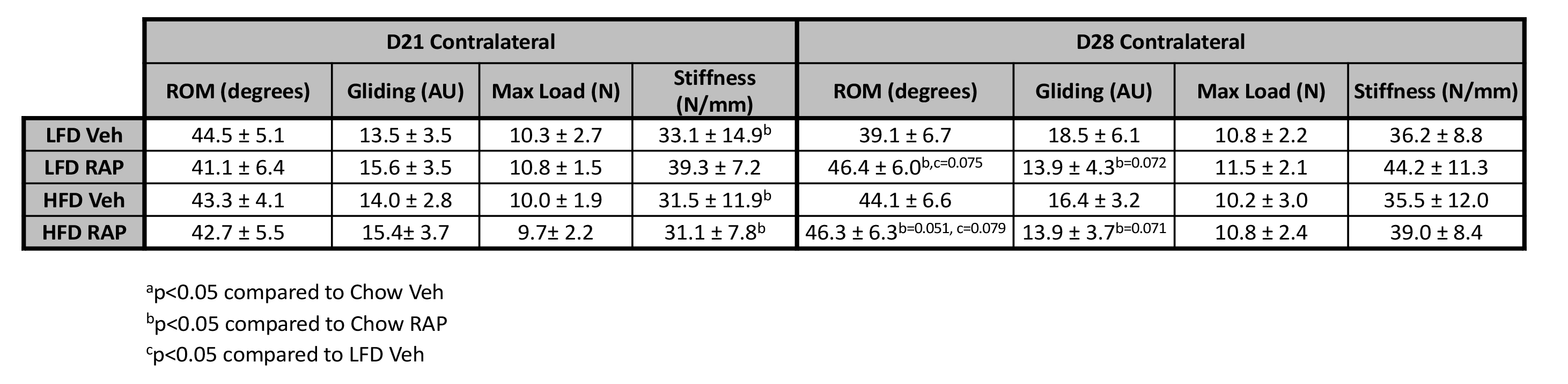

### Supplemental Table 3

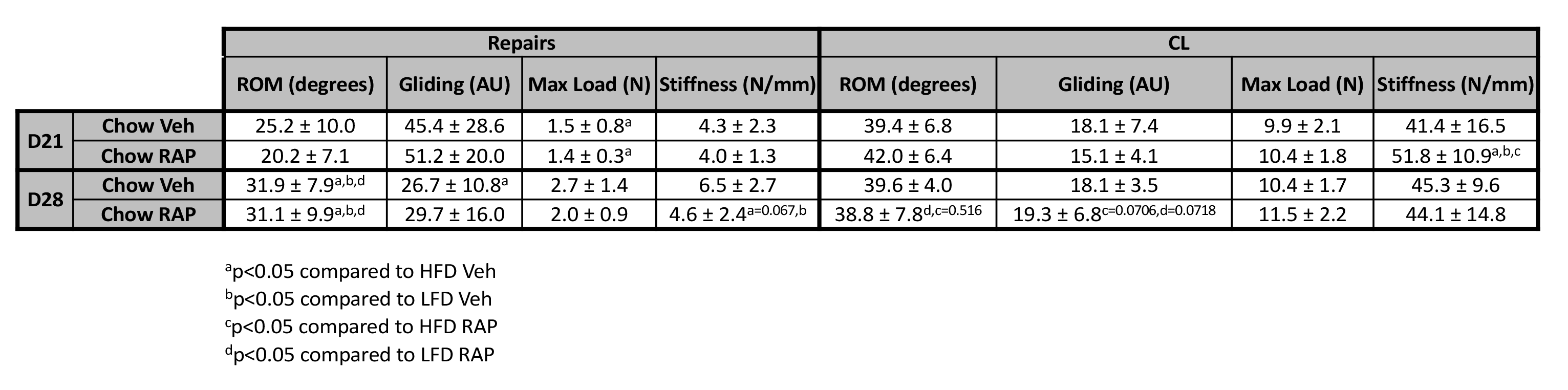
